# The human nose organoid respiratory virus model: an ex-vivo human challenge model to study RSV and SARS-CoV-2 pathogenesis and evaluate therapeutics

**DOI:** 10.1101/2021.07.28.453844

**Authors:** Anubama Rajan, Ashley Morgan Weaver, Gina Marie Aloisio, Joseph Jelinski, Hannah L. Johnson, Susan F. Venable, Trevor McBride, Letisha Aideyan, Felipe-Andrés Piedra, Xunyan Ye, Ernestina Melicoff-Portillo, Malli Rama Kanthi Yerramilli, Xi-Lei Zeng, Michael A Mancini, Fabio Stossi, Anthony W. Maresso, Shalaka A. Kotkar, Mary K. Estes, Sarah Blutt, Vasanthi Avadhanula, Pedro A. Piedra

## Abstract

There is an unmet need for pre-clinical models to understand the pathogenesis of human respiratory viruses; and predict responsiveness to immunotherapies. Airway organoids can serve as an ex-vivo human airway model to study respiratory viral pathogenesis; however, they rely on invasive techniques to obtain patient samples. Here, we report a non-invasive technique to generate human nose organoids (HNOs) as an alternate to biopsy derived organoids. We made air liquid interface (ALI) cultures from HNOs and assessed infection with two major human respiratory viruses, respiratory syncytial virus (RSV) and severe acute respiratory syndrome coronavirus-2 (SARS-CoV-2). Infected HNO-ALI cultures recapitulate aspects of RSV and SARS-CoV-2 infection, including viral shedding, ciliary damage, innate immune responses, and mucus hyper-secretion. Next, we evaluated the feasibility of the HNO-ALI respiratory virus model system to test the efficacy of palivizumab to prevent RSV infection. Palivizumab was administered in the basolateral compartment (circulation) while viral infection occurred in the apical ciliated cells (airways), simulating the events in infants. In our model, palivizumab effectively prevented RSV infection in a concentration dependent manner. Thus, the HNO-ALI model can serve as an alternate to lung organoids to study respiratory viruses and testing therapeutics.

## Introduction

With the onset of the coronavirus disease 2019 (COVID-19) pandemic, it is imperative now more than ever to have pre-clinical, physiologically relevant airway models to evaluate viral infection and potential therapeutics. Airway organoid (AOs) models have quickly become an important investigational tool due to their ability to mimic human respiratory physiology [1]. Previous studies have reported culturing AOs from human induced pluripotent stem (iPS) cells that represent the fetal or developmental stages of the lung [2–6] and the Clevers group recently reported an advanced method for long-term culturing of human lung tissue-derived AOs [7]. Brewington *et al* [8] and Gamage *et al* [9] also modeled nasal epithelial cells using nasal biopsy samples and nasal brushings. However, all the above methods utilize invasive techniques and typically require physicians to obtain lung tissue or bronchoalveolar lavage or nasal brushings from patients, which limit their application to the general researchers and for therapeutic screening. Therefore, a critical need remains for the development of a non-invasive method for generating AOs that can be readily applied to both pediatric and adult populations. Here we report a novel, expandable, ex-vivohuman nose organoid (HNO) model that capitalizes on non-invasive techniques and yet retains the architecture of the respiratory epithelium. Additionally, we have effectively modeled these nasal wash and swab derived HNOs to study the pathogenies of the major pediatric respiratory viral pathogen, respiratory syncytial virus (RSV), and the foremost global respiratory viral pathogen, severe acute respiratory syndrome coronavirus-2 (SARS-CoV-2).

Globally, RSV infection in children <5 years results in 33.1 million cases, 3.2 million hospitalizations, and up to 200,000 deaths, annually [10, 11]. RSV is the major cause of acute lower respiratory tract illness (ALRTI) in children accounting for approximately 20% of all ALRTI [12]. RSV infects almost all children by two years of age, and causes repeated reinfections throughout life [13]. RSV is also a significant cause of respiratory disease morbidity and mortality in older adults, immunocompromised adults, and those with chronic pulmonary disease [14]. Unlike RSV, SARS-CoV-2 pandemic has resulted in a record-breaking global disaster. The rapidly spreading SARS-CoV-2 has caused over 194 million cases and 4.1 million deaths as of July 26, 2021 [15]. Therefore, it is important to develop human model systems to study viral pathology and to test therapeutics against them. In this study, we modeled an HNO-derived air liquid interface (ALI) culture system to grow and study RSV, and SARS-CoV-2 infections. The HNO-ALI cultures were readily infected by contemporaneous RSV strains (RSV/A/Ontario (ON) and RSV/B/Buenos Aires (BA)), and SARS-CoV-2 (WA-1) resulting in epithelial damage reminiscent of human pathology. Cytokine analysis of RSV and SARS-CoV-2 infected HNO epithelium demonstrated cell polarity specific response thus highlighting the importance of using polarized cells to understand the host immune response. Furthermore, this HNO respiratory virus model functions as an ex-vivo human challenge model where we tested the efficacy of palivizumab, a monoclonal antibody used for the prevention of RSV.

## Results

### Development of the Human Nose Organoids (HNOs)

We established six lines of HNOs using stem cells isolated from nasal wash and mid-turbinate swab samples collected from human volunteers adopted from a recently published protocol [7] (Figure 1A). The 3D HNOs formed between 2-3 weeks after initial adaptation of stem cells to specialized growth media conditions (Figure 1B-C). Microscopically, the differentiated HNO-ALI culture system was composed of polarized, pseudostratified airway epithelium containing basal cells (keratin 5 positive, KRT5), secretory club cells SCGB1A1 (CC10 positive), goblet cells (mucin 5AC positive, Muc-5AC), and ciliated cells (acetylated tubulin positive, Ace-tub) (Fig 1D-G). Beating cilia of HNO-ALI were also visible under light microscopy (Movie V1). We performed RNA sequencing of undifferentiated 3D HNOS, and early (21 day) and late differentiated (31 day) HNO-ALI to analyze airway cell specific gene expression pattern. Our data reveals that the HNO-ALI transcriptome was highly enriched for several of airway epithelial cell specific-markers including keratins, dynein, and secretoglobins (Appendix 1); and also showed hallmark of ciliary function as shown by gene set enrichment analysis (GSEA) (Supplemental Figure 1). In summary, our HNOs retained the in-vivo characteristics of human airway epithelium and can also be indefinitely passaged, and frozen for long-term expansion.

**Figure 1:**
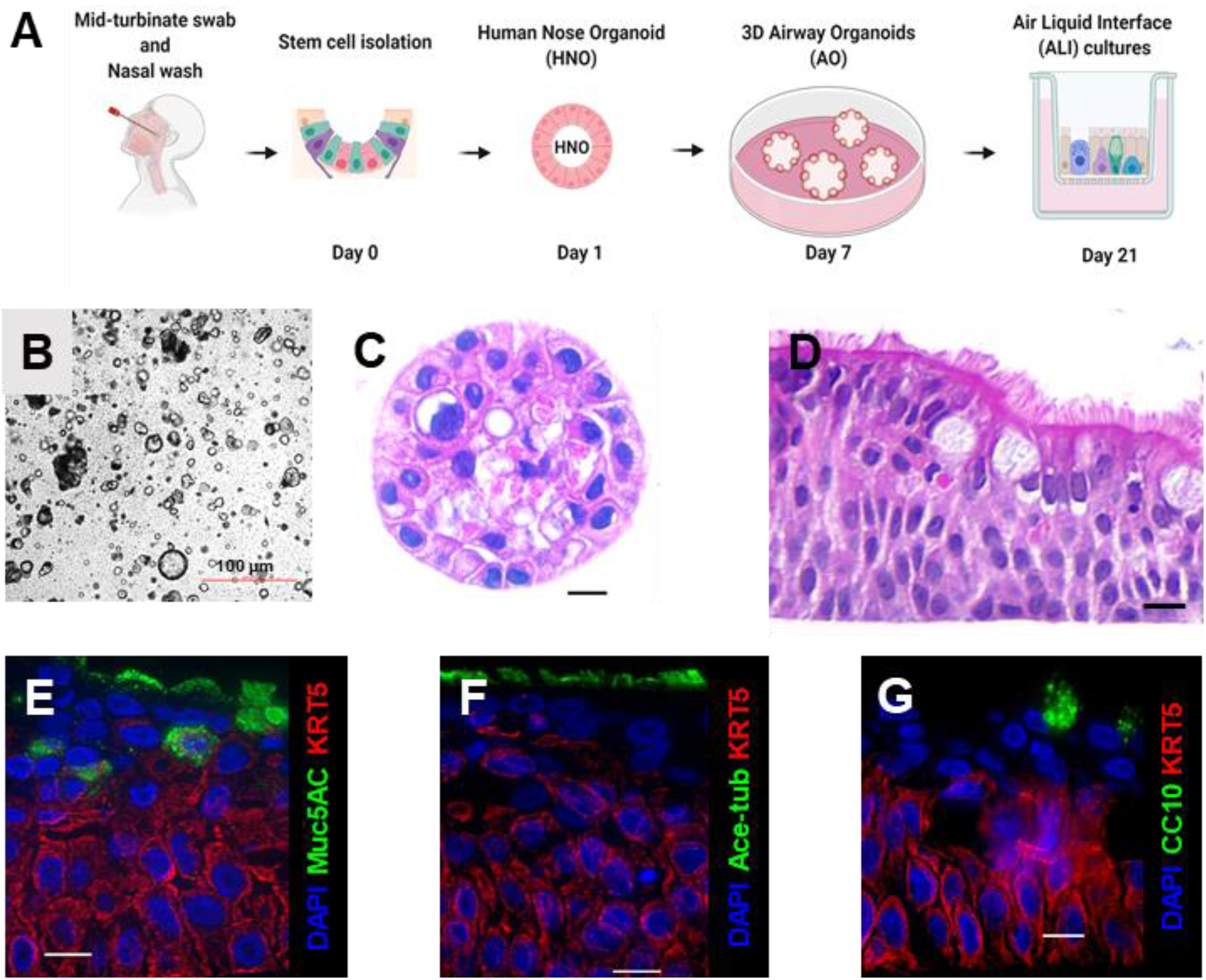
Derivation and characterization of Human Nose Organoids (HNOs). (A) Schematic representation of the workflow for making of HNOs. (B) Bright field image of 3D HNOs in culture. Scale bar equals 100μm. (C) H&E staining of 3D HNOs. (D) H&E staining of pseudostratified airway epithelium of HNO-ALI culture. (E-F). Immunofluorescence of HNOs. Basal cells are labeled by KRT5, in red. DAPI stains each nucleus. (E) Goblet cells are shown in green, labeled by MUC5AC. (F) Ciliated epithelium is shown in green, labeled by Ace-tubulin and (G) Club cells in green, labeled by CC10. Scale bar equals 10μm.

### Replication kinetics of RSV, and SARS-CoV-2 in HNO-ALI system

We tested the ability of the HNO-ALI system to model RSV, and SARS-CoV-2 viral infections using two HNO lines, HNO2 and HNO204. The HNO-ALI system was apically inoculated with two contemporaneous RSV strains, RSV/A/ON and RSV/B/BA, representing RSV/A and RSV/B subtypes. RSV showed robust replication in both the HNO-ALI lines reaching ~ 5X10^7^ RNA copies/ml in the apical compartment by day 5 and reaching steady state over 10 days post inoculation (dpi) (Figure 2A and 2B). The infection of HNO-ALI produced infectious virions as demonstrated by plaque assays on 2, 5, 10 dpi (Figure 2D and E). Live infectious RSV was detected only on the apical side of the HNO-ALI and not on the basolateral compartment, consistent with RSV being predominantly released from the apical cells to the lumen of the HNO-ALI culture. The infection of HNO-ALI with SARS-CoV-2 (WA-1) also showed increases in extracellular viral loads reaching ~ 5X10^8^ RNA copies/ml by day 3 and a plateau at 6-dpi (Figure 2C). In contrast to RSV, SARS-CoV-2 genome was detected both on the apical lumen and the basolateral compartment of the HNO-ALI system suggesting SARS-CoV-2 virions are infecting cells in the apical layer and basolateral layers of the HNO epithelium later into the infection.

**Figure 2:**
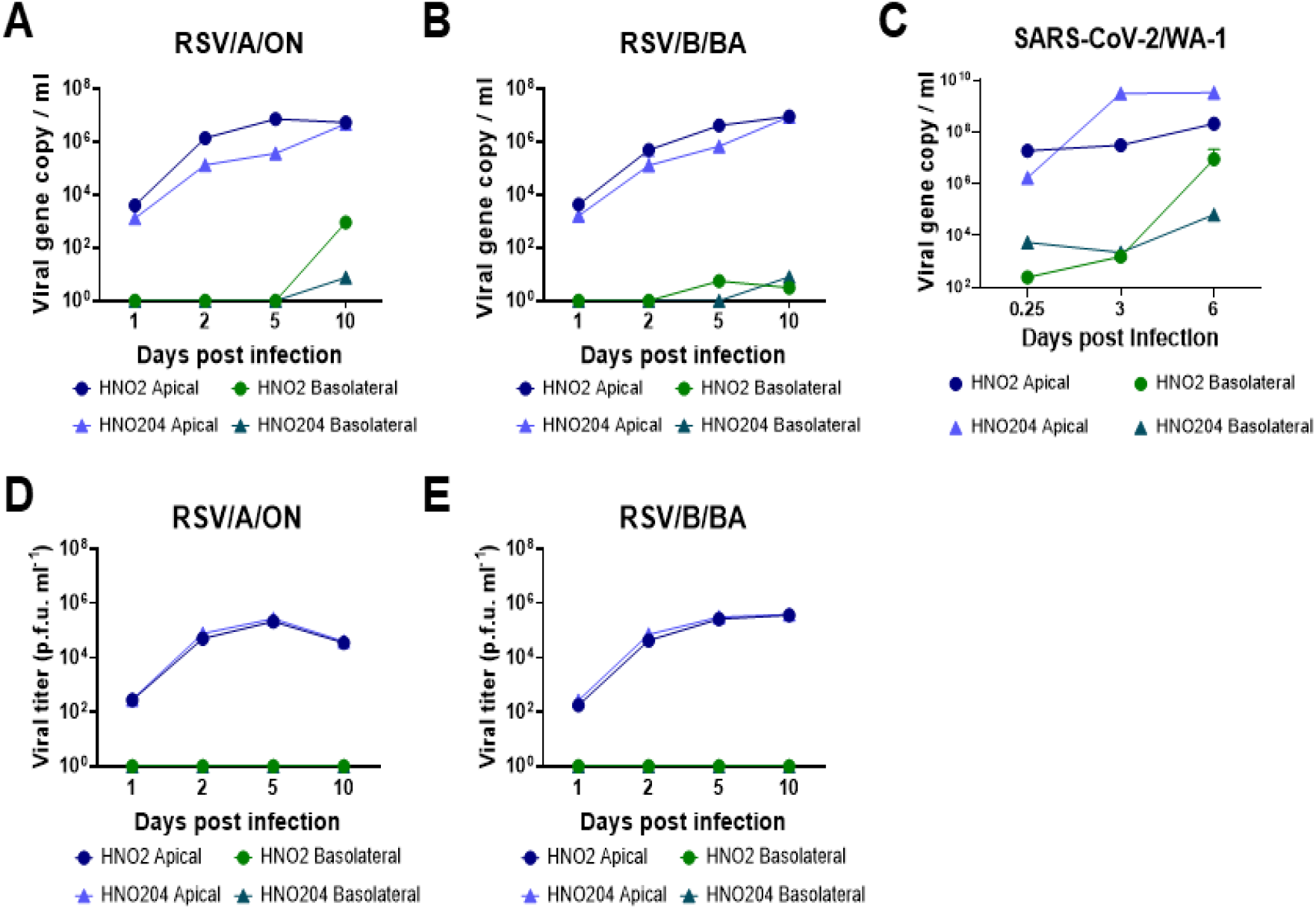
Infection of Human Nose Organoids (HNOs) with RSV and SARS-CoV-2. (A-C) HNO-Air Liquid Interface (ALI) cells were apically infected with RSV/A/Ontario (ON), RSVB/Buenos Aires (BA) at an MOI of 0.01. Apical and basolateral samples were collected at 1-, 2-, 5-, and 10-days post-inoculation (dpi), in two distinct HNO cell lines (HNO2 and HNO204). RNA was isolated from media and copy numbers of RSV N gene RNA were determined using quantitative real time PCR (qRT-PCR). Levels of RSV N gene (copies/ml) from (A) RSV/A/ON at different time points, (B) RSV/A/BA at different time points. (C) HNO-ALI cells were infected with SARS-CoV-2. Apical and basolateral samples were collected at the time of infection, 3- and 6- dpi in two distinct HNO cell lines (HNO2 and HNO204). RNA was isolated from media, SARS-CoV-2 copy numbers of N gene RNA was determined using qRT-PCR. (D-E) Infectious viral titers reported as plaque forming unit (PFU) per ml for RSV/A/ON and RSV/B/BA using a quantitative plaque assay. Data shown were from two individual experiments with two technical replicates per group in each experiment and are represented as mean ± SD.

### Morphological analysis of RSV and SARS-CoV-2 infected HNO-ALI system

Changes in epithelial cell morphology upon RSV and SARS-CoV-2 infection were examined and compared by immunofluorescence labeling of HNO-ALI cultures. Both RSV and SARS-CoV-2 infected HNO-ALI cultures recapitulated in-vivo data including apical shedding of viral infected cells, ciliary damage, and epithelial thinning (Figure 3A-3D). Early infection (6hrs and 1 dpi) was comparable between RSV/A/ON, RSV/B/BA, and SARS-CoV-2, but SARS-CoV-2 infection caused more damage at later time points (6-dpi). Also, RSV infection appeared confined to the apical cell layer. By contrast, SARS-CoV-2 spike protein antigen was detected deeper into the basal cell layer. Using fluorescence threshold analysis to quantify cilia expression (acetylated tubulin or Ace-Tub), we found that SARS-CoV-2 caused significantly higher ciliary damage in comparison to RSV (Figure 3E). The thickness of the epithelium also was significantly lower in SARS-CoV-2 infected cells in comparison to RSV (Figure 3F). The H&E and PAS-AB staining of HNO-ALI further revealed severe extrusion, epithelial thinning and rounding of apical cells in SARS-CoV-2 infected samples (Supplemental Figure 2). In contrast, RSV but not SARS-CoV-2 showed hypersecretion of mucus as quantified by expression of MUC5AC marker (Figure 3C, 3D, 3G and Supplemental Figure 3).

**Figure 3:**
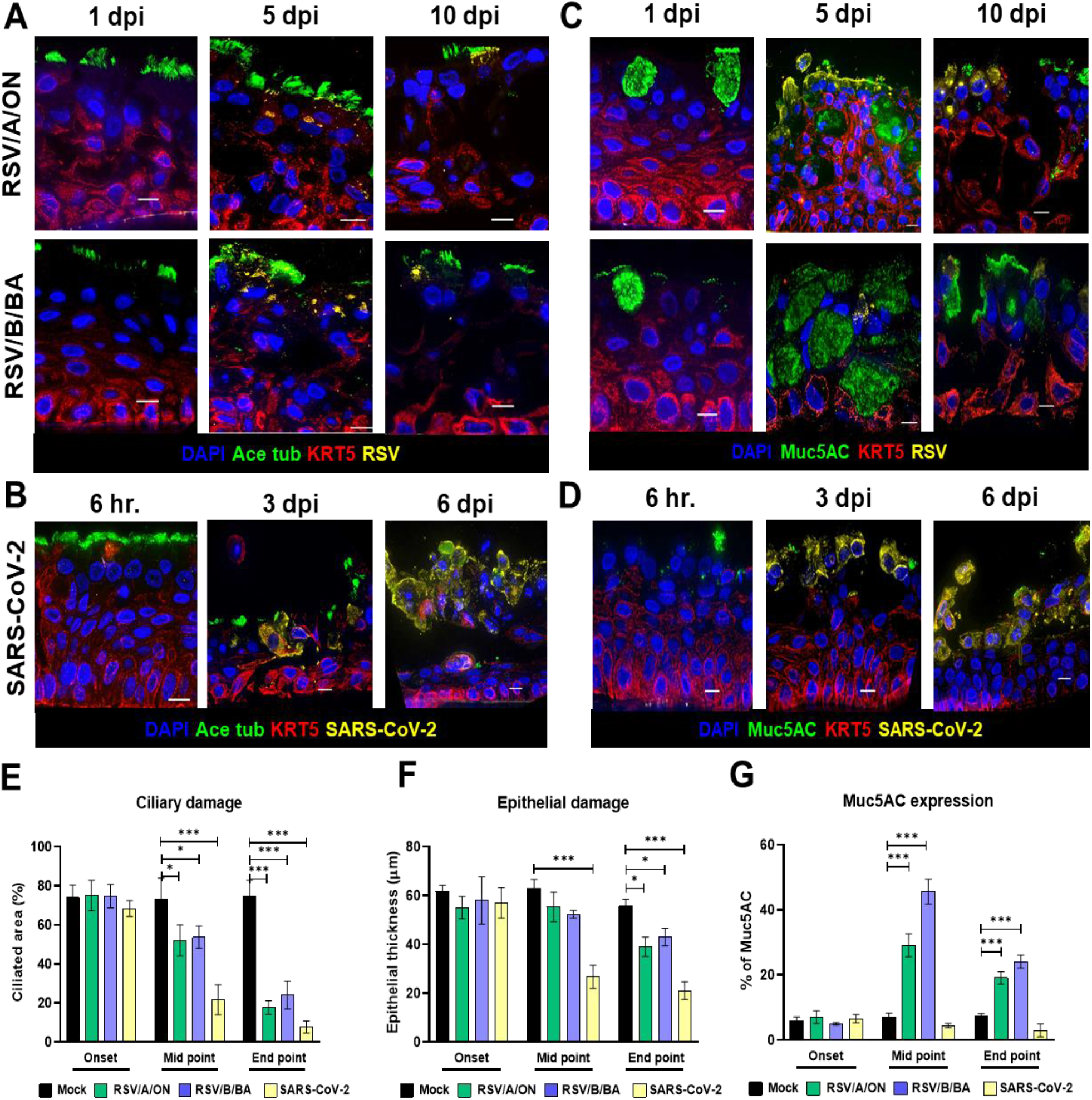
Immunofluorescence and morphological analysis of RSV and SARS-CoV-2 infected Human Nose Organoid-Air Liquid Interface (HNO-ALI) system. (A-D) Representative deconvolved epifluorescence images of HNOs cells showing nuclei (DAPI, blue), and basal cells (KRT5, red). In panel (A) cilia (ace-tubulin, green), and RSV (F-protein, yellow) are shown, in panel (B) cilia (ace-tubulin, green), SARS-CoV-2 (spike protein, yellow, (C) goblet cells (MUC5AC, green), RSV (F-protein, in yellow) and (D) goblet cells (MUC5AC, green), and SARS-CoV-2 (spike protein, yellow). (E-G) Quantification of (E) ciliary damage, (F) epithelial damage and (G) MUC5AC expression of HNO-ALI infected by RSV/A/ON, RSV/B/BA and SARS-CoV2. Data were collected from 10 representative images per group in each experiment and are represented as mean ± SD. (* p=0.05, ** p = 0.01 and *** p = 0.001). Scale bar equals 10μm.

### RSV but not SARS-CoV-2 predominantly induces interferon lambda (IFN-λ1) and IFN-γ inducible chemokine response in HNO-ALI cultures

Characterizing host immune response and identification of biomarkers is crucial towards modeling of RSV and SARS-CoV-2 pathogenesis in these advanced HNO-ALI culture systems. To do this, we performed Luminex cytokine analysis of 29 cytokines/chemokines in HNO2-ALI (Appendix 3). Bronchial and nasal epithelial cells are known to secrete inflammatory cytokines in response to viral infections. We analyzed the levels of inflammatory cytokines induced by HNO-ALI cultures in response to (i) both RSV/A/ON and RSV/B/BA infection at 1-dpi, 2-dpi, 5-dpi, and 10-dpi (Appendix 3) and (ii) SARS-CoV-2 infection at 6hr, 3-dpi, and 6-dpi. RSV infection induced a strong IFN-λ1/IL-29 response in HNO-ALI cultures at 5-dpi and 10-dpi. In striking contrast, HNO2-ALI showed no changes in the levels of IFN-λ1 in response to SARS-CoV-2 infection (Figure 4A and 4G). Notably, for RSV strong inductions were observed for chemokine (C-X-C motif) ligand 10 (IP-10), CXCL9, CXCL11/IP-9 and regulated on activation, normal T cell expressed and secreted (RANTES) from both the apical and basolateral side of the transwells. (Figure 4B – 4E). The C-X-C chemokine ligands are generally upregulated by IFN-gamma (γ) produced from activated T cells and natural killer cells, none of which were present in the HNO-ALI cultures. Next, interleukin-8 (IL-8), a classical biomarker of RSV infection was also detected at 5-dpi and 10-dpi at both the apical and basolateral side of HNO-ALI cultures (Supplemental Table 3) [16]. RSV induced high levels of vascular endothelial growth factor A (VEGF-α) specifically on the basolateral side of HNO-ALI. SARS-CoV-2 infections showed similar trends in the levels of the same cytokines, but the concentration of these cytokines was much lower than RSV (Figure 4I – 4L). In contrast, increase in CXCL10 (IP-10) by ~100-fold was measured on the basolateral side of SARS-CoV-2 infected samples at 6-dpi, which has been reported as a biomarker of COVID-19 disease severity (Figure 4H) [17].

**Figure 4:**
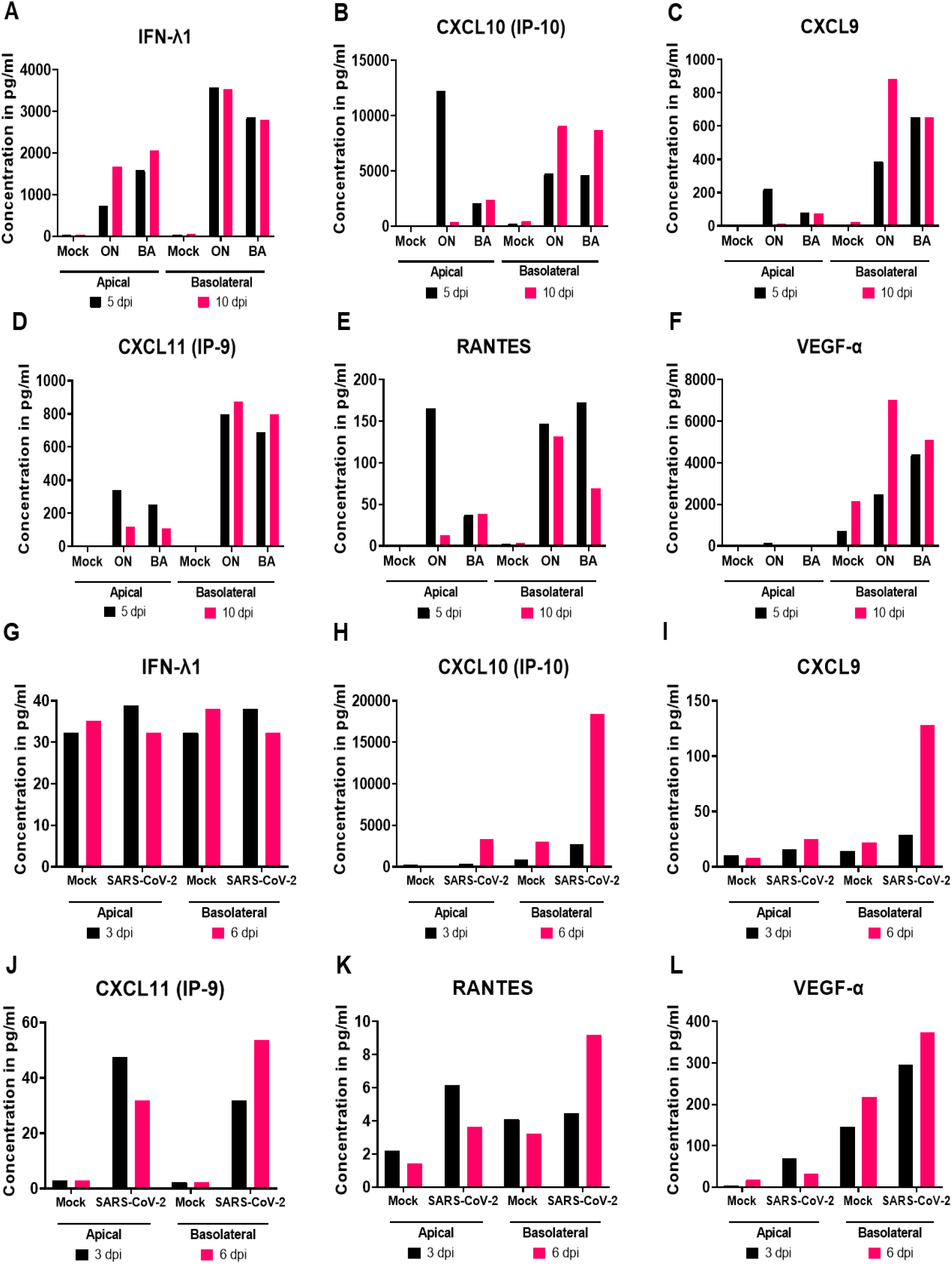
Immune cytokine/chemokine profile of Human Nose Organoids (HNOs) infected with Respiratory Syncytial Virus (RSV) and severe acute respiratory syndrome coronavirus 2 (SARS-CoV-2). Cells were infected with RSV/A/Ontario (ON), RSVB/Buenos Aires (BA) at a multiplicity of infection (MOI) of 0.01. The cultured supernatants were harvested from mock, RSV and severe acute respiratory syndrome coronavirus-2 (SARS-CoV-2) infected cells. Profiles of extracellular cytokines and chemokines released in the supernatants were determined by the Multiplex-Luminex cytokine assay. (A-F) Levels of cytokines and chemokines released from HNO2 cells infected with contemporaneous RSV strains RSV/A/ON and RSV/B/BA. (G-L) Levels of cytokines and chemokines released from HNO2 cells infected with SARS-CoV-2 (WA-1 strain).

Increase in matrix metalloproteinase 9 (MMP-9) was reported in COVID-19 patients with respiratory failure [18] and during RSV infection [19]. However, we did not detect significant changes in the levels of MMP-9 or MMP-7 to either RSV or SARS-CoV-2 (Appendix 3). Perhaps, this could be due to the inhibitory effect of TIMP-1 on MMPs. Additionally, for both RSV and SARS-CoV-2, we did not detect any changes in levels of some pulmonary fibrosis biomarkers such as transforming growth factor beta (TGF-β), fibroblast growth factor (FGF), granulocyte colony-stimulating factor (G-CSF), and granulocyte-macrophage colony-stimulating factor (GM-CSF) [20]. On the other hand, VEGF was elevated and has been associated with disease severity in idiopathic pulmonary fibrosis [20]. Though measured in low amounts (less than 100 pg/ml), we noticed an increase in levels of proinflammatory immune mediators such as IL-6, and IL-1α for both RSV and SARS-CoV-2 as seen in clinical disease (Appendix 3). In contrast, only for RSV and at the basolateral side, monocyte chemoattractant protein 1 (MCP-1), MIP-1β, and tumor necrosis factor-α (TNF-α) levels increased at 5-dpi. Eotaxin, IL-1β, and MCP-3 were both increased at apical and basolateral side at low levels in response to RSV infection (Appendix 3).

### Ex-vivo human challenge respiratory virus model in the HNO-ALI system

Although multiple pre-clinical animal models are available to recapitulate some pathognomonic aspects of RSV and SARS-CoV-2 infection, they do not faithfully represent the physiology of human airway epithelium [21–26]. Additionally, there are advanced iPS and lung tissue derived AO models where the disease can be modeled but they require access to clinical samples to establish these culture systems [7, 27, 28]. Thus, there is an unmet and urgent need for physiologically relevant, yet easily accessible pre-clinical human airway models for respiratory viral diseases to test the efficacy of therapeutics. To address this, we tested the ability of the HNO-ALI system to act as an ex-vivo human challenge respiratory virus model to test the efficacy of a known therapeutic monoclonal antibody (mAb) to RSV infection. Palivizumab (Synagis^®^), is a neutralizing mAb targeted against the F-glycoprotein of RSV that prevents RSV–cell fusion and hence reduces RSV replication [29]. We introduced the palivizumab mAb in the basolateral compartment and monitored its neutralizing capacity on the apical lumen mimicking the neutralizing effects of mAb in circulation on the virus exposed airway epithelium (Supplemental Figure 3B). The palivizumab concentrations used were in the biological range shortly after intravenous injection (640μg/ml) or prior to the next administration dose (80μg/ml) [30].

HNO2-ALI cultures pre-incubated with palivizumab at 640μg/ml suppressed RSV/A/Tracy replication up until 2-dpi. Thereafter, RSV replication resumed at 4-dpi although at reduced levels as compared to the no palivizumab control, and replication reached a peak at 6-dpi, and finally plateaued at 8-dpi (Figure 5A). This demonstrated both the efficacy of palivizumab to reduce infection in the apical ciliated cells and also suggested a decline in bioavailability of palivizumab to persistently prevent RSV replication at later time points. In a subsequent experiment, HNO-ALI cultures were pre-incubated with palivizumab prior to the virus inoculation and given a second dose at 4-dpi. In this experimental setup, RSV/A/Tracy failed to replicate in HNO-ALI culture at both 640μg/ml and 80μg/ml of palivizumab (Figure 5B). Nonetheless, palivizumab resistant strain-RSV/Tracy^P-Mab R^ [31] readily replicated in palivizumab pre-treated HNO-ALI cultures demonstrating the specificity of palivizumab in preventing RSV infection (Figure 5C). Furthermore, we analyzed the cytokines expressed by palivizumab pre-incubated-HNOs that were inoculated with RSV. Palivizumab pre-treatment with either the 80 or 640 μg/ml dose, followed with second dose not only prevented RSV replication (Figure 5B and 5E) but also abolished the virus-induced inflammatory cytokines that would have been released into the apical lumen and basolateral compartment (Figure 5G – 5K).

**Figure 5:**
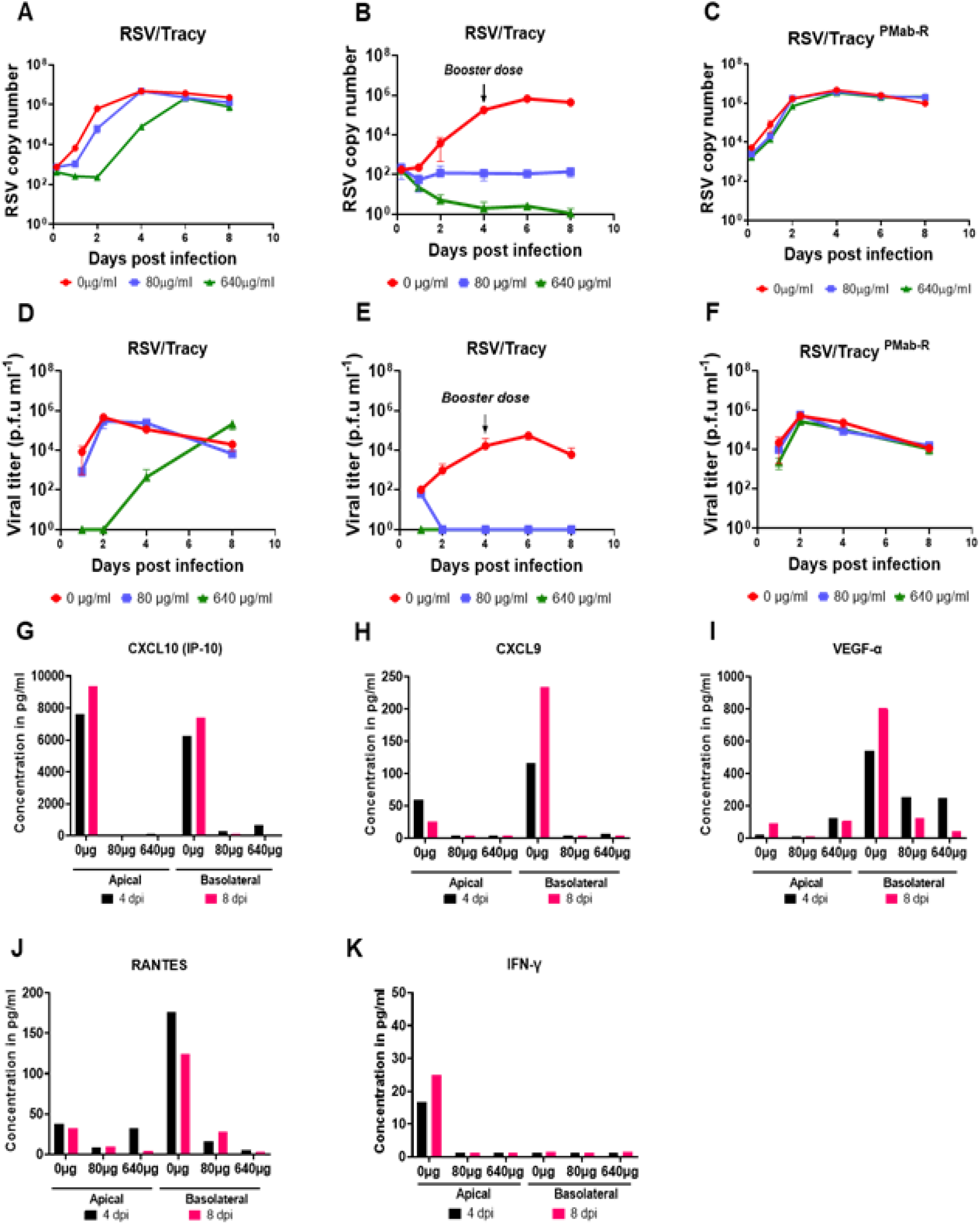
Immunoprophylaxis treatment for Respiratory Syncytial Virus (RSV) infection in Human Nose Organoids (HNOs). (A-C) shows viral copy number of RSV/A/Tracy after treatment with palivizumab of 0μg/ml, 80μg/ml, and 640μg/ml at 1, 2, 4, 6, and 8 dpi were measured. (A) demonstrates only an initial but not sustained decrease in RSV copy number with a single dose of Palivizumab pre-treatment. (B) shows a significant reduction in RSV replication across the infection period with the addition of a “booster” dose of Palivizumab at day 4, in a dose-dependent manner. (C) exhibits specificity of Palivizumab pre-treatment, as strain RSV/A/Tracy^-Mab-R^ is resistant to the antibody, and hence showed no decrease in infection. (D-F) Quantification of infective viral particles using plaque assay for the experiments described above. (G-K) shows the levels of inflammatory cytokines released by RSV-infected HNOs under Palivizumab treatment conditions.

## Discussion

Airway organoids are three-dimensional airway culture systems that were first developed in 1993 with self-organizing 3D structures [32]. To date, there are only a handful of AO models that are produced using invasive lung biopsy or bronchoalveolar lavage (BAL) patient-donor materials [4, 6, 7, 33–39]. In this study for the first time, we describe a non-invasive, reproducible and a reliable approach to establish human nose organoids (HNOs) that allows for long-term expansion. Earlier studies used invasive nasal brushing samples or biopsy samples that are traditionally obtained from human subjects undergoing bronchoscopy procedures. In our protocol, we used human nasal wash and mid-turbinate swab samples from healthy volunteers to generate nasal organoids. The ease in obtaining the nasal wash/mid-turbinate swab samples facilitates our non-invasive approach into the general adult population as well as the vulnerable pediatric population. These nasal wash and mid-turbinate samples can either be self-collected or through a trained laboratory technician. A notable difference between our HNO-ALI culture and other ALI is the richness and stratification of airway epithelium [8, 36–39]. While nasal epithelial cells reported by *Muller et al, Brewington et al, and Liu et al* can be grown in ALI conditions they are predominantly represented as a single monolayer of cells. In contrast, our HNO-ALI system derived from HNOs exhibits a complex pseudostratified epithelium interlaced with basal, goblet, club, and ciliated cells with spontaneous synchronization of beating cilia (Figure 1D-G and Movie V1). In short, this newly developed HNO model offers an elegant solution to develop in-vitro human respiratory airway models that can be used across several basic-science laboratories to model respiratory diseases.

RSV infections cause millions of hospitalizations every year [10] and almost all children are infected with RSV by two years of age with repeated reinfections occurring throughout life [13]. Another most important pathogen in the current scenario is SARS-CoV-2, the causative organism of COVID-19. AO systems are known for the robustness to model respiratory viral infections such as SARS-CoV-2 [35, 40], influenza virus [34], enterovirus [41], and other respiratory diseases [1]. We tested the usefulness of our HNOs as ex-vivo culture system for studying two major human respiratory viruses: RSV, and SARS-CoV-2. Traditional cell culture models including immortalized and primary airway epithelial cell lines, viral inoculum is introduced in the cell culture media. This limits the exposure of air to airway epithelium, which is a critical component for physiological relevance. Additionally, most of the existing 3D airway organoid models have the apical side facing inward and microinjections are needed to establish infection. In our HNO-ALI model, the apical side of the epithelium is both accessible and air exposed, thus making it both physiologically relevant and highly suitable for studying respiratory viral infections. Here, we report that the HNO-ALI system readily supports the replication and growth of RSV and SARS-CoV-2 (Figure 2A-2C) and the newly released virions from these cultures are infectious (Figure 2D, and 2E). Using immunohistochemistry, we also showed that the apical ciliated epithelial cells are indeed the main target for RSV (Figure 3A). These results are consistent with previously published studies on AOs and RSV where apical infection of ciliated cells was observed [7, 42–44]. Although we did not detect any RSV infection of basal cells, in contrast to our observation, a previous study has noted that basal cells if exposed to RSV are susceptible to RSV infection suggesting a pathogenic mechanism for additional airway damage in individuals with chronic lung disease [45]. Next, in our study we observed that only RSV induced a robust increase in expression of MUC5AC (Figure 3C) and this was comparable to the *Persson et al* study where they have also reported expansion of MUC5AC secretory cells in response to RSV infection [45]. Unlike RSV, SARS-CoV-2 induced severe damage to the ciliary epithelium without an increase in mucus secretion (Figure 3B, and 3D).

We performed a thorough characterization of host immune response to RSV and SARS-CoV-2 infection in HNO-ALI system. Although RSV infection predominantly occurred at the apical surface, we measured higher levels of cytokine response to viral infection on the basolateral side suggesting translocation of infection signals from the apical side to the basolateral compartment mimicking physiological relevance. We also observed a strong type-III interferon (IFN-λ1/IL-29) response for RSV and not SARS-CoV-2 suggesting stark differences in the role of the epithelial cells in initiating the early innate and antiviral immune signaling. It was interesting to observe that, the increase in IFN-λ1/IL-29 responses to RSV at 5-dpi was associated with a reduced or stalled viral replication beyond 5-dpi as noted by plateauing of viral copy number measured by RT-PCR (Figure 2A, 2B and 4A) or infectious virus (Figure 2D, 2E and 4D) measured by a plaque assay. Additionally, a previous study has shown association of IFN-λ1/IL-29 and increase in MUC5AC secretory cells during RSV infection [45]. Consistent with these findings our data also demonstrates both increase in IFN-λ1 and MUC5AC expression in response to RSV infection in HNO-ALI cultures (Figure 3B, 3D and 4A). However, future studies are needed to clarify the mechanisms of hypersecretion of MUC5AC induced by RSV in HNO-ALI system.

IFN-γ inducible cytokine/chemokine response (IP-9, CXCL-9 and RANTES) and increase in VEGF-α was noted to both RSV and SARS-CoV-2 infection (Figure 4). This data is in accordance with clinical findings in patients from previously published studies, where high levels of IFN-γ and CXCL10 (IP-10) were seen for RSV [46–48]. IP-10, a biomarker for both RSV and COVID-19 severity was detected in the infected HNO2 cultures. In contrast, a study conducted by *Rijsbergen et al* showed no increase in IP-10 in response to RSV infection in nasal epithelial cells [44]. An important remark: previous studies using airway epithelial culture system have shown and supported the growth of RSV and SARS-CoV-2 only up to 4-dpi [7, 28, 40]. However, in our system, we extended the viral infection for RSV beyond 8 days and for SARS-CoV-2 beyond 6 days. This allows for long-term study of viral infections. Thus, we hope that this HNO-derived airway epithelial model will greatly enhance the understanding of RSV and SARS-CoV-2 pathogenesis.

Currently, there is an unmet need for an easily available preclinical human model to study RSV and SARS-CoV-2 infections that recapitulates the human experience. While animal models of infection have provided major insights, they have also misdirected our understanding of viral pathogenesis and prevention [25]. Also, the cost, safety and ethical issues associated with human challenge models limit their use and access [49]. Our ex-vivo HNO-ALI model is an alternative to the human challenge model. In our study, HNO-ALI cultures were pre-treated with palivizumab to model immunoprophylaxis treatment to prevent RSV infection. A single dose of palivizumab pre-treatment of HNO-ALI culture was only partially effective in reducing RSV replication (1-dpi at 80μg/ml; and up to 2-dpi at 640μg/ml). Nonetheless, when a second dose of palivizumab was administered at 4-dpi, RSV was unable to replicate in HNO-ALI system and the HNOs did not produce inflammatory cytokines in response to RSV infection demonstrating the immunoprophylactic protection against RSV infection and disease. Indeed, our HNO-ALI system more closely resembled the human experience where therapeutic mAb or polyclonal antibodies are administered intramuscularly or intravenously, respectively, to get into the blood circulation and provide protection of the airways against RSV infection [50–52]. Thus, these HNOs will provide a more precise human milieu and can function as a pre-clinical human model to investigate promising therapeutics while recapitulating the complex interactions between the drug, the virus, and the airway cells.

We are of the opinion that, at the current stage, the HNO-ALI system remains at a highly reductionist level and has the potential for improvements. This includes advancements of HNO-ALI system with 1) addition of immune cells, 2) endothelial cells to further mimic the complex physiology of organs and 3) furthermore genetic knock-ins and knockouts can be made to study the role of specific host components in microbial infections and human diseases. With these future advancements, the HNOs can be optimized to develop next-generation in-vivo human airway models and used as a valuable tool to evaluate pathogenesis, therapeutics, and vaccine candidates for major global respiratory viral pathogens. In addition, the HNOs retain the genetic background of the individual, thus allowing the possibility to screen drugs for cancer therapeutics, genetic-disease modeling, and development of personalized medicine.

## Methods

### Obtaining nasal wash/ nasal swab samples

The samples were collected under the Institutional Review Board (IRB) of the Baylor College of Medicine (BCM), Houston, Texas, USA with written informed consent. Self-collected nasal wash samples were obtained by instilling 3 ml of saline into each nostril and collecting the fluid into a sterile cup. A paired mid-turbinate nasal swab from the same volunteer was also obtained using a flocked swab. The paired nasal wash and nasal swab samples were mixed and stored on ice until further processed.

### Establishing Human Nose Organoids (HNOs)

The generation of nasal wash and nasal swab derived HNOs was based on the published protocol [7]. The nasal wash along with the flocked swab was spun at 80g for 5 minutes at 4⁰C. Supernatant was carefully removed and sheared using a 29-gauge insulin syringe to break up mucus (if any). Digestion medium [10ml AO medium + 10mg Collagenase (Sigma C9407) +100μl Fungizone (Amphotericin B)] was added and the falcon tube was kept on a rocker for 30-60 minutes at 37⁰C. After digestion, the nasal swab was discarded, and fetal bovine serum (FBS) was added to inactivate collagenase. The above solution was sheared using a syringe, strained through a 100μm strainer, and spun at 80g for 5 minutes at 4⁰C. The supernatant was removed, and the cell pellet was washed twice with Wash Medium (96 ml Advanced DMEM/F12 + 1 ml Glutamax 100x + 1 ml HEPES 1M + 1ml Pen/Strep + 1ml Fungizone) and spun at 80g for 5 minutes at 4⁰C. Finally, the Wash Medium was removed, and the cell pellet was suspended in Matrigel^®^ and plated onto a 24 well plate and incubated for 10 minutes at 37⁰C. Once Matrigel^®^ had solidified, 500μl of AO medium with Penicillin Streptomycin Amphotericin (PSA) was added to each well and the plate was transferred into 37⁰C incubator. AO medium was replaced every 4 days and passaged every other week at 1:2 ratio (wells) for expansion.

### Generation of HNO-Air Liquid Interface (ALI) cultures

The mature 3D HNOs were enzymatically and mechanically sheared to make ALI cultures using our previous method adopted from enteroid monolayer technology and conditions utilized for growing human bronchial epithelial cells [33, 53–56]. Clear transwells (Corning Costar, Catalog # 3470) were pre-coated with 100μl of Bovine Type I collagen at 30μg/ml (Gibco, Catalog # A1064401) and placed in an incubator for 1.5 hour at 37⁰C. HNO cultured in AO medium for 7 days were dissociated using 0.5mM EDTA and spun at 300g for 5 minutes at 4⁰C. Single cells were obtained by adding 0.05% Trypsin/0.5mM EDTA (Invitrogen, Catalog # 25300054) for 4 minutes at 37⁰C. Trypsin was inactivated by addition of AO medium containing 10% FBS. The HNOs were dissociated vigorously using pipette tips, passed through 40μm strainer (Falcon, Catalog # 352340) and pelleted at 400g at room temperature for 5 minutes to generate single cells. The pellet was resuspended in AO medium containing 10μM Y-27632 + Epidermal Growth Factor (EGF) (Peprotech-AF-100-15). The collagen coating from transwells was removed, washed with phosphate buffered saline (PBS) and the single cells were added at a seeding density of 3 x 105 cells/well. 750μl of AO medium + EGF containing 10μM Y-27632 (Sigma, Catalog # Y-0503) was added into the lower compartment of the transwells. After 4 days, confluent monolayers are cultured in air-exposed conditions using differentiation medium (PneumaCult-ALI medium from STEMCELL Technologies) in lower compartments for Transwell until 21 days.

### Viral infection, PCR, and plaque assays

An overview of the infection method is provided in Supplemental Figure 3A. The differentiated HNO-ALI cells were apically infected with RSV/A/USA/BCM813013/2013(ON) (RSV/A/ON), RSV/B/USA/BCM80171/2010(BA) (RSV/B/BA), SARS-CoV-2 [Isolate USA-WA1/2020, obtained from Biodefense and Emerging Infectious resources (BEI)] at a multiplicity of infection 0.01 (30μl/well). All work with SARS-CoV-2 was performed in a Class II Biosafety Cabinet in the BSL-3 high-containment facility at BCM. For mock infection, AO-differentiation media (30 μl/well) alone was added. The inoculated plates were incubated for 1.5 hours 37 °C with 5% CO2. At the end of incubation, the inoculum was removed from the HNO-ALI system and left air-exposed for the defined period of infection. At the respective time points, the apical side of the transwells was washed twice with 150μl of AO differentiation media and mixed with equal volume of 15% glycerol Iscoves media. On the basolateral side, 300μl of the media was removed and mixed with equal volume of 15% glycerol Iscoves media. The obtained samples were used for detection of viral RNA, infectious virions, and host cytokines. Samples were snap frozen and stored at −80⁰C. The viral RNA was extracted using Mini Viral RNA Kit (Qiagen Sciences) in an automated QIAcube platform according to the manufacturer instructions [57]. Viral RNA was detected using real time polymerase chain reaction (RT-PCR) with primers targeting the nucleocapsid gene for RSV, SARS-CoV-2 as previously described [57, 58]. RSV titer was measured using semi-quantitative plaque assay as previous described [59].

### Immunohistochemistry (IHC), and Immunofluorescence labeling (IF)

HNO-ALI cultures were fixed in image-iT™ Fixative Solution (4% formaldehyde) [Catalog number: FB002] for 15 minutes followed by dehydration in ethanol series (30%, 50%, 70% and 90%, each 30 minutes at room temperature or overnight at 4⁰C). The transwells membranes were then subjected to paraffin embedding, and sectioning. Standard H&E (hematoxylin and eosin) and PAS/AB (Periodic acid–Schiff/ Alcian-Blue) staining was performed. For immunofluorescent labeling, the sections were deparaffinized in Histo-Clear (Electron Microscopy Science, 64111-01), followed by washes in an alcohol sequence (100>100>95>65 %). The slides were then rehydrated and exposed for heat-induced antigen retrieval in 10 mM Sodium Citrate buffer pH 6 or 20 mins at sub-boiling temperature [60]. The sections were rinsed in water and blocked for 60 min in 1% bovine serum albumin (BSA) in Tris buffered saline with Tween20 (TBS-T) (blocking buffer). The sections were incubated overnight at 4°C with the following primary antibodies, 0.25 μg/ml keratin 5 (KRT5; BioLegend, Catalog number: 905503), 0.2μg/ml SCGB1A1 (CC10; Santa Cruz, Catalog number: sc-365992 AF488), 0.2μg/ml acetylated alpha tubulin (Santa Cruz, sc-23950 AF488), 0.2μg/ml Mucin 5AC (Invitrogen, Catalog number: 45M1), goat anti-RSV IgG antibody (Abcam, Catalog number: ab20745) and rabbit anti-SARS-CoV-2 S1 IgG Antibody (Sino biologicals, Catalog number: 40150R007; BEI Resources, Catalog number: NR10361, 1:2000 dilution of the antiserum). Primary antibodies were washed three times in TBS+0.05 % Tween for 10 min each, incubated with secondary antibodies from Invitrogen, for 1hour at room temperature, washed twice with TBS-T, and stained with 4′,6-diamidino-2-phenylindole (DAPI), washed twice with PBS, and mounted in VECTASHIELD Plus Antifade mounting media (Vector Laboratories, Burlingame, CA, H-1900). The slides were stored at 4°C until imaging.

### Immunofluorescence image quantification and analysis

High quality/high resolution automated imaging was performed on a Cytiva DV Live epifluorescence image restoration microscope using an Olympus Plan Apo N 60X/1.42 NA objective and a 1.9k x 1.9k pco. EDGE sCMOS_5.5 camera with a 1024×1024 FOV. The filter sets used were DAPI (390/18 excitation, 435/48 emission), FITC (475/28 excitation, 523/36 emission), TRITC (542/27 excitation, 594/45 emission), and CY5 (632/22 excitation, 676/34 emission). Z stacks (0.25μm) of the whole section (~10μm) were acquired before applying a conservative restorative algorithm for quantitative image deconvolution using SoftWorx v7.0. Max intensity projections were used for image analysis and processed using ImageJ/Fiji. Between 6 and 10 slides were imaged per treatment/biological replicate and analyzed for IF studies. Every experiment was performed in two technical transwell replicates and repeated for minimum of two biological replicates. For visualization, RSV/SARS-CoV-2 spots were enhanced by histogram stretching across treatments, post image acquisition in Fiji. The average area of ciliated epithelium was quantified by cell counts with acetylated-tubulin in Fiji at the beginning of each infection (day 1 for RSV, 6hr for SARS-CoV-2), the midpoint (day 5 for RSV, 3 days for SARS-CoV-2), and the endpoint of infection assays (day 10 for RSV and day 6 for SARS-CoV-2). Quantification of amount of epithelial damage by RSV-A, and SARS-CoV-2 was measured by calculating the average thickness of the epithelium in μm. Percentage of goblet cells (MUC5AC labeled area) relative to DAPI after infection with RSV-A, RSV-B and SARS-CoV-2 was measured using the formula “μm^2^ MUC5AC label / μm^2^ of epithelium (DAPI)” at the beginning of each infection (day 1 for RSV, 6hr for SARS-CoV-2), the midpoint (day 5 for RSV, 3 days for SARS-CoV-2), and the endpoint of infection assays (day 10 for RSV and day 6 for SARS-CoV-2). GraphPad Prism 9.0 was used to construct graphs and perform statistical tests.

### Multiplex Luminex Immunoassays

Cytokines and chemokines secreted by HNO-ALI cells were measured and analyzed using Milliplex cytokine/chemokine magnetic bead panel (Millipore) according to the manufacturer’s instructions. The kits used in this study include 1) Milliplex Human Cytokine Panel with Eotaxin/CCL11, FGF-2, G-CSF, GM-CSF, IL-1a, IL-1b, IL-6, IL-8/CXCL8, IL-17E/IL-25, IP-10/CXCL10, MCP-1, MCP-3, MIG, MIP1a, MIP1b, RANTES/CCL5, TNFa, VEGF-A, IL-33, TRAIL, TSLP, TAC/CXCL11, IL-29, BAFF and HMGB1, 2) TGFb1 Singleplex kit, 3) Milliplex Human MMP Panel 2 with MMP9 and MMP7 and 4) Milliplex Human TIMP Panel 2 with TIMP1. Data were obtained with Luminex xPONENT for MAGPIX, version 4.2 Build 1324 and analyzed with MILLIPLEX Analyst version 5.1.0.0 standard Build 10/27/2012.

### Immunoprophylaxis model of HNO-ALI

To establish and test the feasibility of HNO-ALI system to serve as an ex-vivo human airway challenge model, we developed an immunoprophylaxis system using palivizumab antibodies to prevent RSV infection. An overview of the immunoprophylaxis model is summarized in Supplemental Figure 3B. The differentiated HNO-ALI cells were apically infected with RSV/A/USA/BCM-Tracy/1989 (GA1) and RSV-Tracy resistant to Palivizumab at a multiplicity of infection 0.01 (30μl/well). In our model, palivizumab was introduced on the basolateral compartment either 30 minutes prior to RSV infection or at both 30 minutes prior to RSV infection and a second dose at 4-dpi. Palivizumab was administered on the basolateral side of HNO-ALI system at 80μg/ml; and 640μg/ml.

### Statistical analysis

A statistical significance was determined by two-way ANOVA. Tukey’s multiple comparison tests are performed using GraphPad Prism version 7.0 Windows (Graph Pad Software, San Diego, CA, www.graphpad.com). Differences in the means are considered significant at p≤0.05 with specific p values detailed in the figure legends.

## Acknowledgements

The model figures were created with BioRender.com.

These studies were supported in part by grant U19 AI144297 that is as part of the U19 program (Genomic Centers of Infectious Diseases-GCID) from the National Institutes of Health (NIH). Cytokine analysis through Luminex platform was supported by Public Health Service Grant P30DK056338, funding to the Texas Medical center Digestive Disease Center. Imaging for this project was supported by the Integrated Microscopy Core at Baylor College of Medicine and the Center for Advanced Microscopy and Image Informatics (CAMII) with funding from NIH (DK56338, CA125123, ES030285), and CPRIT (RP150578, RP170719), the Dan L. Duncan Comprehensive Cancer Center, and the John S. Dunn Gulf Coast Consortium for Chemical Genomics. Funders had no role in design of experiment, data collection or interpretation of data.

---

**Supplemental Figure 1:**
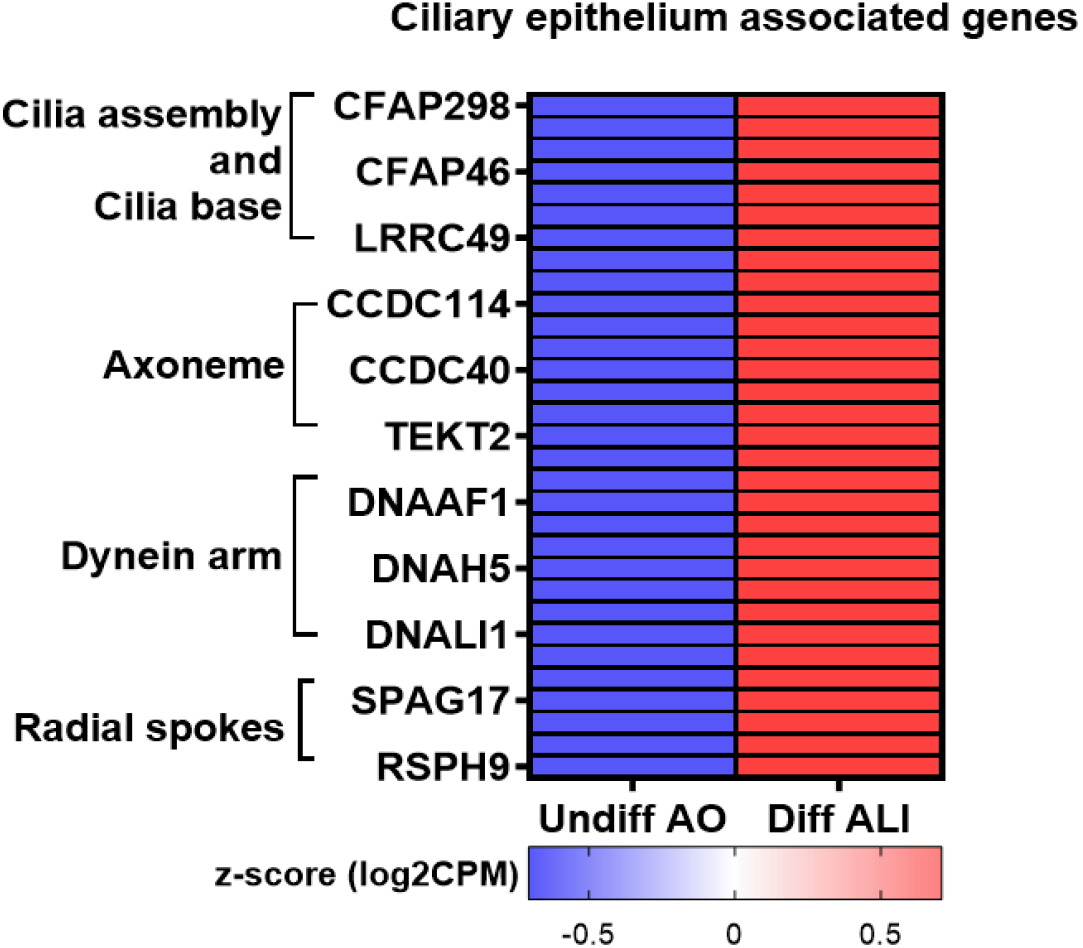
List of ciliary epithelia associated genes in Human Nose Organoids (HNOs). Heatmap summarizes a list of expression of ciliary associated genes in undifferentiated 3D AO cultures and 21-day differentiated ALI cultures. This is suggestive of effective differentiation process inducing ciliogenesis and functional ciliary movements in HNO-ALI cultures.

**Supplemental Figure 2:**
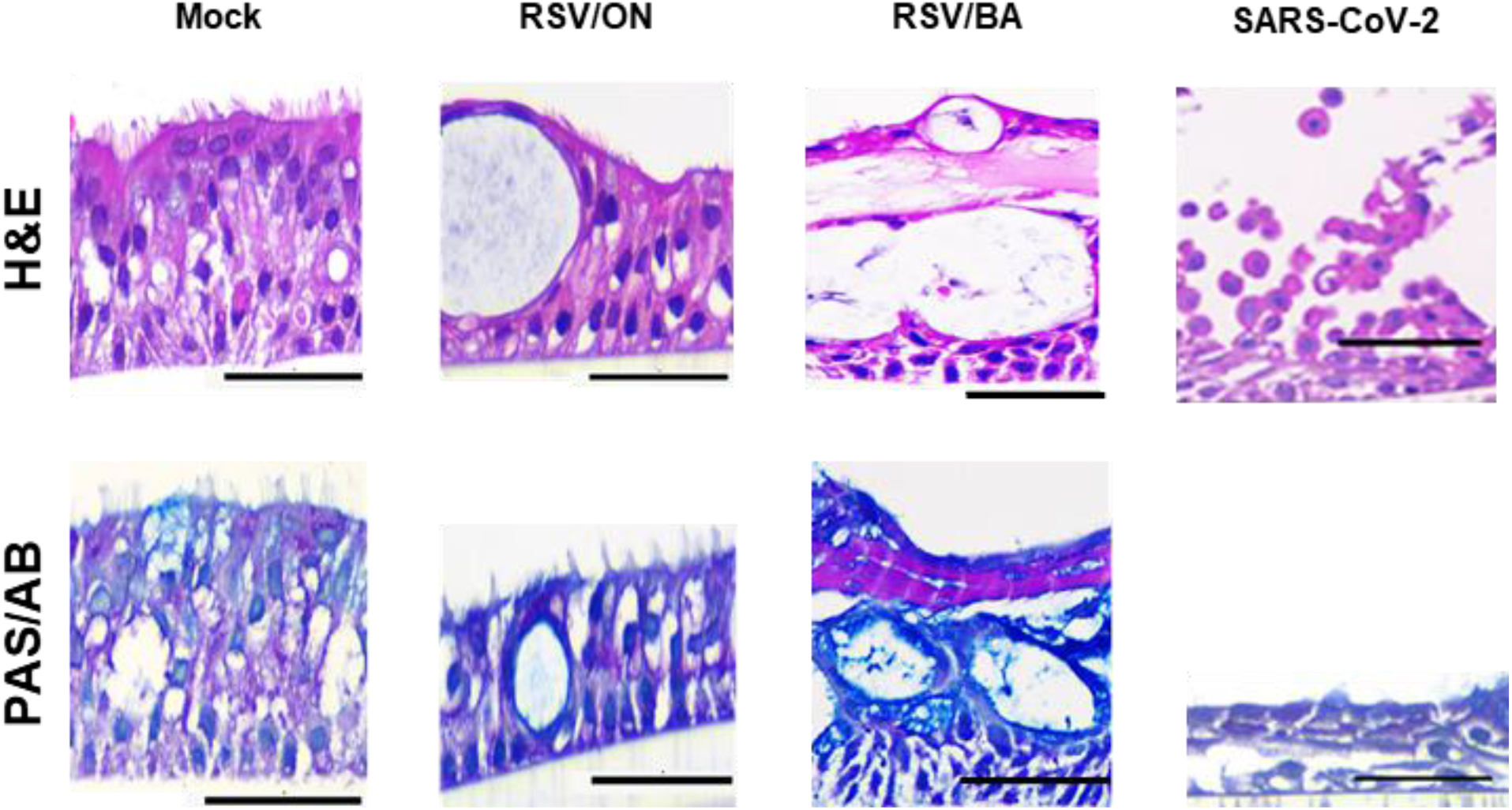
Histological characterization of Human Nose Organoids (HNOs) infected with respiratory viruses: RSV/A/Ontario (ON), RSV/B/Buenos Aires (BA) and Severe Acute Respiratory Syndrome Coronavirus-2 (SARS-CoV-2).

**Supplemental Figure 3:**
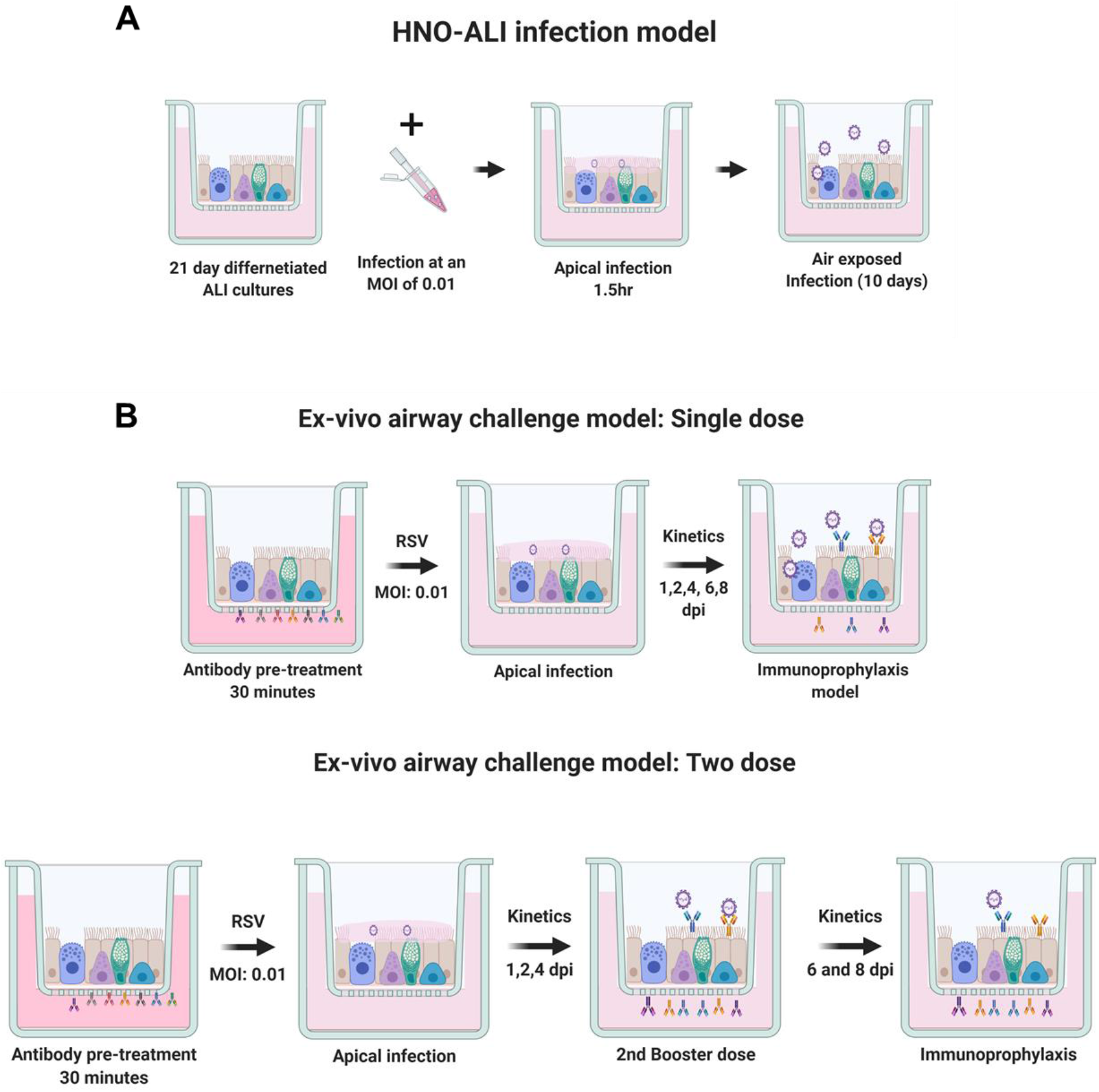
Human Nose Organoid-Air Liquid Interface (HNO-ALI) infection and airway challenge model.

**Supplemental Table 1:**
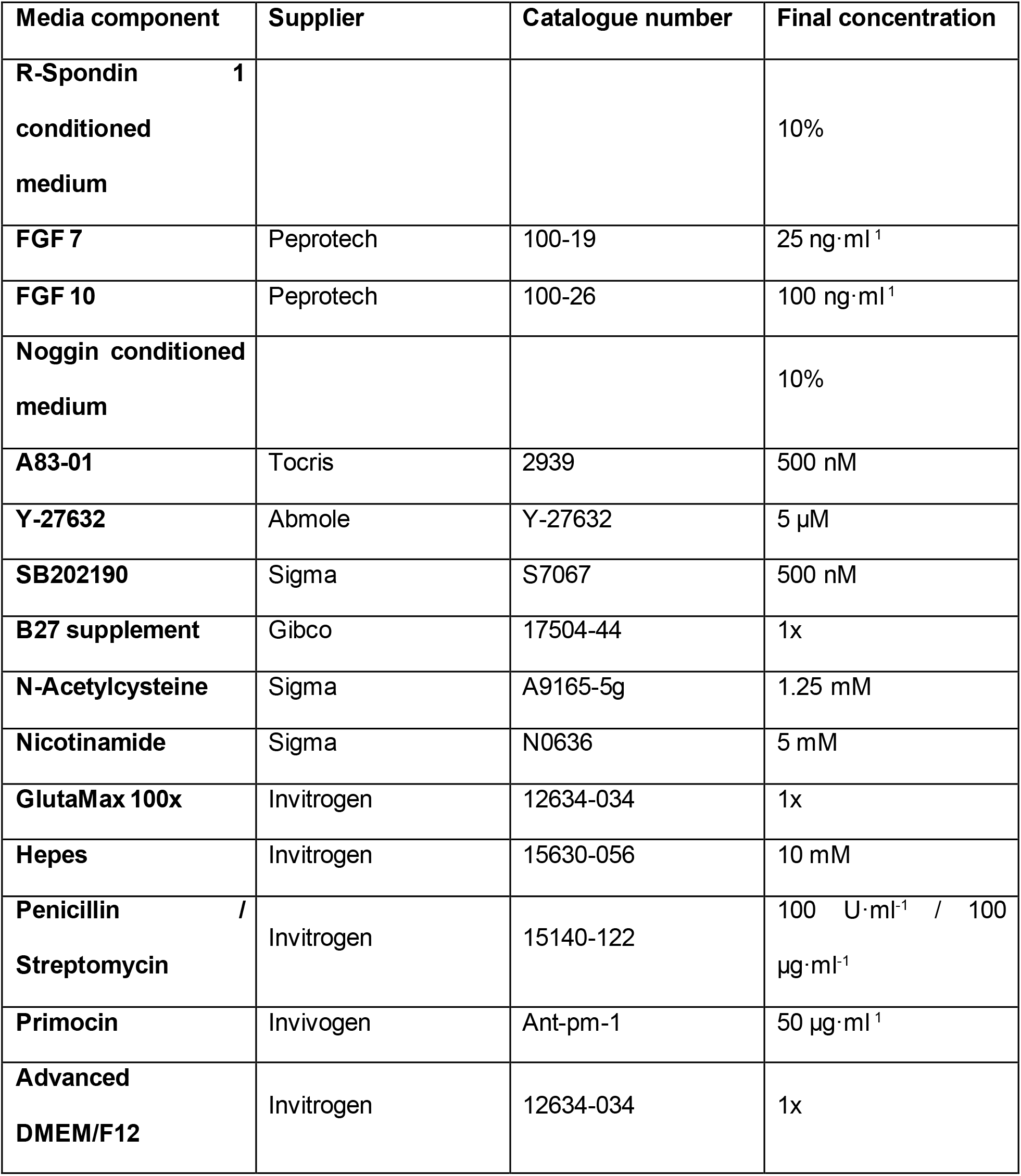
Airway Organoid (AO) medium composition.

## References

1. van der Vaart, J. and H. Clevers, Airway organoids as models of human disease. Journal of Internal Medicine. n/a(n/a).

2. Tadokoro, T., et al., IL-6/STAT3 promotes regeneration of airway ciliated cells from basalstem cells. Proceedings of the National Academy of Sciences, 2014. 111(35): p. E3641.

3. Tadokoro, T., et al., BMP signaling and cellular dynamics during regeneration of airway epithelium from basalprogenitors. Development, 2016. 143(5): p. 764–73.

4. Dye, B.R., et al., In vitro generation of human pluripotent stem cell derived lung organoids. eLife, 2015. 4: p. e05098.

5. Chen, Y.-W., et al., A three-dimensionalmodelof human lung development and disease from pluripotent stem cells. Nature Cell Biology, 2017. 19(5): p. 542–549.

6. Huang, S.X., et al., Efficient generation of lung and airway epithelial cells from human pluripotent stem cells. Nat Biotechnol, 2014. 32(1): p. 84–91.

7. Sachs, N., et al., Long-term expanding human airway organoids for disease modeling. Embo j, 2019. 38(4).

8. Brewington, J.J., et al., Detection of CFTR function and modulation in primary human nasalcell spheroids. J Cyst Fibros, 2018. 17(1): p. 26–33.

9. Gamage, A.M., et al., Infection of human Nasal Epithelial Cells with SARS-CoV-2 and a 382-nt deletion isolate lacking ORF8 reveals similar viral kinetics and host transcriptional profiles. PLoS Pathog, 2020. 16(12): p. e1009130.

10. Nair, H., et al., Global burden of acute lower respiratory infections due to respiratory syncytial virus in young children: a systematic review and meta-analysis. Lancet, 2010. 375(9725): p. 1545–55.

11. Shi, T., et al., Global, regional, and national disease burden estimates of acute lower respiratory infections due to respiratory syncytial virus in young children in 2015: a systematic review and modelling study. The Lancet, 2017. 390(10098): p. 946–958.

12. Welliver, T.P., et al., Severe human lower respiratory tract illness caused by respiratory syncytial virus and influenza virus is characterized by the absence of pulmonary cytotoxic lymphocyte responses. J Infect Dis, 2007. 195(8): p. 1126–36.

13. Hall, C.B., The burgeoning burden of respiratory syncytial virus among children. Infect Disord Drug Targets, 2012. 12(2): p. 92–7.

14. Falsey, A.R., et al., Respiratory syncytial virus infection in elderly and high-risk adults. N Engl J Med, 2005. 352(17): p. 1749–59.

15. Dong, E., H. Du, and L. Gardner, An interactive web-based dashboard to track COVID-19 in real time. The Lancet Infectious Diseases, 2020. 20(5): p. 533–534.

16. Vázquez, Y., et al., Cytokines in the Respiratory Airway as Biomarkers of Severity and Prognosis for Respiratory Syncytial Virus Infection: An Update. Frontiers in Immunology, 2019. 10(1154).

17. Chen, Y., et al., IP-10 and MCP-1 as biomarkers associated with disease severity of COVID-19. Molecular Medicine, 2020. 26(1): p. 97.

18. Ueland, T., et al., Distinct and early increase in circulating MMP-9 in COVID-19 patients with respiratory failure. J Infect, 2020. 81(3): p. e41–e43.

19. Dabo, A.J., et al., Matrix Metalloproteinase 9 Exerts Antiviral Activity against Respiratory Syncytial Virus. PLOS ONE, 2015. 10(8): p. e0135970.

20. Guiot, J., et al., Blood Biomarkers in Idiopathic Pulmonary Fibrosis. Lung, 2017. 195(3): p. 273–280.

21. Zhang, A.J., et al., Severe Acute Respiratory Syndrome Coronavirus 2 Infects and Damages the Mature and Immature Olfactory Sensory Neurons of Hamsters. Clinical Infectious Diseases, 2020.

22. Williamson, B.N., et al., Clinical benefit of remdesivir in rhesus macaques infected with SARS-CoV-2. Nature, 2020. 585(7824): p. 273–276.

23. Richard, M., et al., SARS-CoV-2 is transmitted via contact and via the air between ferrets. Nat Commun, 2020. 11(1): p. 3496.

24. Johansen, M.D., et al., Animal and translational models of SARS-CoV-2 infection and COVID-19. Mucosal Immunology, 2020. 13(6): p. 877–891.

25. Taylor, G., Animal models of respiratory syncytial virus infection. Vaccine, 2017. 35(3): p. 469–480.

26. Bem, R.A., J.B. Domachowske, and H.F. Rosenberg, Animal models of human respiratory syncytial virus disease. American Journal of Physiology-Lung Cellular and Molecular Physiology, 2011. 301(2): p. L148–L156.

27. Mulay, A., et al., SARS-CoV-2 infection of primary human lung epithelium for COVID-19 modeling and drug discovery. bioRxiv, 2020.

28. Lamers, M.M., et al., An organoid-derived bronchioalveolar model for SARS-CoV-2 infection of human alveolar type II-like cells. The EMBO Journal, 2021. 40(5): p. e105912.

29. Updated guidance for palivizumab prophylaxis among infants and young children at increased risk of hospitalization for respiratory syncytial virus infection. Pediatrics, 2014. 134(2): p. e620–38.

30. Subramanian, K.N., et al., Safety, tolerance and pharmacokinetics of a humanized monoclonal antibody to respiratory syncytial virus in premature infants and infants with bronchopulmonary dysplasia. MEDI-493 Study Group. Pediatr Infect Dis J, 1998. 17(2): p. 110–5.

31. Gilbert, B.E., et al., Respiratory syncytial virus fusion nanoparticle vaccine immune responses target multiple neutralizing epitopes that contribute to protection against wild-type and palivizumab-resistant mutant virus challenge. Vaccine, 2018. 36(52): p. 8069–8078.

32. Benali, R., et al., Tubule formation by human surface respiratory epithelial cells cultured in a three-dimensional collagen lattice. Am J Physiol, 1993. 264(2 Pt 1): p. L183–92.

33. Jiang, D., N. Schaefer, and H.W. Chu, Air-Liquid Interface Culture of Human and Mouse Airway Epithelial Cells. Methods Mol Biol, 2018. 1809: p. 91–109.

34. Zhou, J., et al., Differentiated human airway organoids to assess infectivity of emerging influenza virus. Proceedings of the National Academy of Sciences, 2018. 115(26): p. 6822.

35. Gamage, A.M., et al., Infection of human Nasal Epithelial Cells with SARS-CoV-2 and a 382-nt deletion isolate lacking ORF8 reveals similar viral kinetics and host transcriptional profiles. PLOS Pathogens, 2020. 16(12): p. e1009130.

36. Cao, H., et al., Testing gene therapy vectors in human primary nasal epithelial cultures. Molecular Therapy - Methods & Clinical Development, 2015. 2.

37. Charles, D.D., et al., Development of a Novel ex vivo Nasal Epithelial Cell Model Supporting Colonization With Human Nasal Microbiota. Frontiers in Cellular and Infection Microbiology, 2019. 9(165).

38. Leung, C., et al., Structural and functional variations in human bronchial epithelial cells cultured in air-liquid interface using different growth media. American Journal of Physiology-Lung Cellular and Molecular Physiology, 2020. 318(5): p. L1063–L1073.

39. Au - Müller, L., et al., Culturing of Human Nasal Epithelial Cells at the Air Liquid Interface. JoVE, 2013(80): p. e50646.

40. Lamers, M.M., et al., SARS-CoV-2 productively infects human gut enterocytes. Science, 2020. 369(6499): p. 50.

41. Freeman, M.C., et al., Respiratory and intestinal epithelial cells exhibit differential susceptibility and innate immune responses to contemporary EV-D68 isolates. bioRxiv, 2021: p. 2021.01.05.425225.

42. Zhang, L., et al., Respiratory syncytial virus infection of human airway epithelial cells is polarized, specific to ciliated cells, and without obvious cytopathology. J Virol, 2002. 76(11): p. 5654–66.

43. Villenave, R., et al., In vitro modeling of respiratory syncytial virus infection of pediatric bronchial epithelium, the primary target of infection in vivo. Proceedings of the National Academy of Sciences of the United States of America, 2012. 109(13): p. 5040–5045.

44. Rijsbergen, L.C., et al., Human Respiratory Syncytial Virus Subgroup A and B Infections in Nasal, Bronchial, Small-Airway, and Organoid-Derived Respiratory Cultures. mSphere, 2021. 6(3).

45. Persson, B.D., et al., Respiratory Syncytial Virus Can Infect Basal Cells and Alter Human Airway Epithelial Differentiation. PLOS ONE, 2014. 9(7): p. e102368.

46. Nicholson, E.G., et al., Robust Cytokine and Chemokine Response in Nasopharyngeal Secretions: Association With Decreased Severity in Children With Physician Diagnosed Bronchiolitis. The Journal of Infectious Diseases, 2016. 214(4): p. 649–655.

47. Sung, R.Y., et al., A comparison of cytokine responses in respiratory syncytial virus and influenza A infections in infants. Eur J Pediatr, 2001. 160(2): p. 117–22.

48. Rajan, A., et al., Multiple RSV strains infecting HEp-2 and A549 cells reveal cell line-dependent differences in resistance to RSV infection. bioRxiv, 2021: p. 2021.06.15.448622.

49. Lambkin-Williams, R., et al., The human viral challenge model: accelerating the evaluation of respiratory antivirals, vaccines and novel diagnostics. Respiratory Research, 2018. 19(1): p. 123.

50. Groothuis, J.R., et al., Prophylactic Administration of Respiratory Syncytial Virus Immune Globulin to High-Risk Infants and Young Children. New England Journal of Medicine, 1993. 329(21): p. 1524–1530.

51. Siber, G.R., et al., Comparison of antibody concentrations and protective activity of respiratory syncytial virus immune globulin and conventional immune globulin. J Infect Dis, 1994. 169(6): p. 1368–73.

52. Group, T.I.-R.S., Palivizumab, a Humanized Respiratory Syncytial Virus Monoclonal Antibody, Reduces Hospitalization From Respiratory Syncytial Virus Infection in High-risk Infants. Pediatrics, 1998. 102(3): p. 531–537.

53. Neuberger, T., et al., Use of primary cultures of human bronchial epithelial cells isolated from cystic fibrosis patients for the pre-clinical testing of CFTR modulators. Methods Mol Biol, 2011. 741: p. 39–54.

54. Poole, N.M., A. Rajan, and A.W. Maresso, Human Intestinal Enteroids for the Study of Bacterial Adherence, Invasion, and Translocation. Curr Protoc Microbiol, 2018. 50(1): p. e55.

55. Saxena, K., et al., Human Intestinal Enteroids: a New Model To Study Human Rotavirus Infection, Host Restriction, and Pathophysiology. Journal of Virology, 2016. 90(1): p. 43–56.

56. Rajan, A., et al., Novel Segment- and Host-Specific Patterns of *Enteroaggregative Escherichia coli* Adherence to Human Intestinal Enteroids. mBio, 2018. 9(1): p. e02419–17.

57. Avadhanula, V., et al., Infection With Novel Respiratory Syncytial Virus Genotype Ontario (ON1) in Adult Hematopoietic Cell Transplant Recipients, Texas, 2011-2013. The Journal of Infectious Diseases, 2015. 211(4): p. 582–589.

58. Avadhanula, V., et al., Viral load of SARS-CoV-2 in adults during the first and second wave of COVID-19 pandemic in Houston, TX: the potential of the super-spreader. J Infect Dis, 2021.

59. Englund, J.A., et al., Rapid diagnosis of respiratory syncytial virus infections in immunocompromised adults. J Clin Microbiol, 1996. 34(7): p. 1649–53.

60. Shi, S.R., et al., Antigen retrieval technique utilizing citrate buffer or urea solution for immunohistochemicaldemonstration of androgen receptor in formalin-fixed paraffin sections. J Histochem Cytochem, 1993. 41(11): p. 1599–604.

